# Measuring the strength of spatial sorting

**DOI:** 10.1101/2025.01.17.633577

**Authors:** Nikunj Goel, Mattheau S. Comerford, Anastasia Bernat, Scott P. Egan, Thomas E. Juenger, Timothy H. Keitt

## Abstract

Empirical work suggests that spatial sorting—a mechanism of evolutionary change fueled by spatial assortment of phenotypes—may lead to phenotypic shifts on ecological timescales. However, we currently lack theoretical tools to measure the strength of spatial sorting, as we do to measure the strength of natural selection. To address this gap, we present a quantitative genetics model and identify an evolutionary parameter in the model to measure the strength of spatial sorting. This parameter, the standardized sorting gradient, is structurally akin to the standardized selection gradient, commonly used to measure the strength of natural selection. To show the utility of our approach, we analyzed wing-morphology data of soapberry bugs (*Jadera haematoloma*) recolonizing flooded habitats extirpated by a hurricane. We found that the estimated strength of spatial sorting ranked in the top ten percentile of standardized selection gradients documented in the scientific literature. Our results underscore that, like natural selection, spatial sorting, too, can yield rapid evolution after extreme events.

## Introduction

Extreme climatic events are becoming common due to rising carbon emissions and concomitant planetary warming (Solow 2015, Stott 2016). These climatic perturbations may impose strong selection on populations, fueling rapid evolution (Grant et al. 2017). Indeed, many studies have documented strong selection in response to extreme events (Bumpus 1899, Boag and Grant 1981, Grant and Grant 1993, Brown and Brown 1998, Donihue et al. 2018). These studies underscore a common theme—an extreme event yields non-random mortality wherein some individuals live past the event because of a genetically determined trait that confers a survival advantage. When these survivors reproduce, the mean phenotype of the population changes, resulting in evolution by natural selection (Darwin and Wallace 1858).

Recent empirical work suggests that extreme events can also produce drastic phenotypic changes by a relatively lesser-known evolutionary mechanism that does not necessarily require non-random survival or reproduction (Comerford et al. 2023). Extreme events can extirpate some local populations in a patchy landscape, creating empty patches (van Bergen et al. 2020). These local extinctions create an opportunity for non-random immigration from nearby extant populations, where a subset of individuals colonize the empty patches due to a genetically heritable trait that confers a dispersal advantage. When these immigrants, who possess a trait that confers them a dispersal advantage, mate with each other, the mean phenotype in hitherto empty patches changes relative to the nearby extant populations. This mechanism of evolution, driven by spatial assortment of phenotypes in a moving frame of reference (rather than temporal assortment in a static frame of reference, as is the case in natural selection), is referred to as spatial sorting (Shine et al. 2011; see Figs. S1 and S2 for a more detailed discussion).

Although a growing number of studies have documented evolution by spatial sorting in both natural (MacKay and Lamb 1979, Peroni 1994, Hill et al. 1999, Hanski et al. 2004, Phillips et al. 2006, Duckworth 2008, Forsman et al. 2011, Lombaert et al. 2014) and experimental settings (Williams et al. 2016, Ochocki and Miller 2017, Szűcs et al. 2017, Weiss-Lehman et al. 2017, Usui and Angert 2024), we lack theoretical tools to measure the strength of spatial sorting from phenotypic data, especially after an extreme event. This gap in the literature stems from two reasons. First, extreme events are hard to predict, making it challenging to plan longitudinal studies to quantify their evolutionary impacts (Grant et al. 2017). Second, the theory of spatial sorting is still in its infancy (Bolnick and Otto 2013, Novak and Kollár 2017, Phillips and Perkins 2019, Peischl and Gilbert 2020). Unlike the Breeder’s equation (Lush 1937) and its extensions (Falconer 1960, Lande and Arnold 1983, Walsh and Lynch 2018) that provide quantitative tools to measure the strength of selection using standardized selection gradients, we do not have a similar quantitative genetics theory that may allow biologists to study phenotypic evolution by spatial sorting. Addressing this knowledge gap is crucial for providing quantitative evidence that spatial sorting can be strong enough to influence ecology, producing novel spatial eco-evolutionary dynamics (Phillips et al. 2006, Ochocki and Miller 2017, Weiss-Lehman et al. 2017, Miller et al. 2020).

This paper has two major goals. First, we derive a quantitative genetics model of spatial sorting to quantify evolutionary change during colonization of empty habitats. This model is analogous to the Breeder’s equation, but it differs in how evolutionary change is quantified. Unlike the Breeder’s equation, which quantifies evolution in a static frame of reference within a population, our quantitative genetics model quantifies evolution by spatial sorting in a moving frame of reference to capture the spatial assortment of phenotypes across ecological patches (Shine et al. 2011; also see Figs. S1 and S2). This quantitative genetics model serves several functions, such as facilitating a conceptual understanding of how spatial sorting drives the evolution of quantitative traits and providing a standardized measure to quantify the strength of spatial sorting, which we refer to as the standardized sorting gradient. Based on the knowledge gained from the model, biologists can plan what data to collect, how to analyze it, and how to interpret the results in the broader context of evolutionary theory. Second, to demonstrate the utility of the quantitative genetics model, we analyze wing morphology data of red-shoulder soapberry bugs (*Jadera haematoloma*; hereafter soapberry bug) collected during a multi-year monitoring experiment (2016-2020) in the Greater Houston area, which overlapped with Hurricane Harvey in 2017 (Comerford et al. 2023). Specifically, using the survey data, we estimate the standardized sorting gradient to quantify the strength of spatial sorting resulting from the phenotype-dependent recolonization of habitats that were extirpated by the hurricane. We also perform experimental crosses to evaluate if the wing morphology has a genetic basis and to test if the non-random recolonization could have resulted in an evolutionary response.

### Quantitative genetics model of spatial sorting

Consider a patchy landscape consisting of randomly distributed vacant and occupied patches. The occupied patches are collectively inhabited by a large population of 𝑁 parents. We denote the trait value of the *i*th parent as 𝜙_𝑖_, which determines individuals’ dispersal capacity and lifetime reproductive success. We denote the mean trait value of the parents by 𝜙^#^. The bar is a shorthand for the mean. Next, some parents disperse from occupied patches to vacant patches. To track the dispersing parents, we define an indicator variable 𝑉_𝑖_, which is one if the 𝑖th parent disperses to a vacant patch and zero otherwise. Following dispersal, the 𝑖th parent produces 𝑊_𝑖_ offspring, such that the mean trait value of offspring, 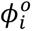, is 𝛿_𝑖_ more than the trait value of the parent.

Based on these notations, the mean trait value of offspring in the vacant patches is

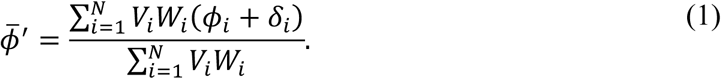

This equation is obtained by adding the trait value of offspring in the vacant patches and then dividing this sum by the number of offspring in the vacant patches. We can re-express the above equation compactly as

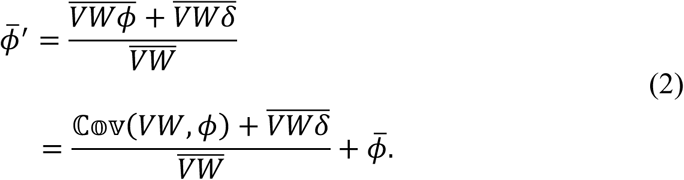

By re-arranging the terms in equation (2), we find that, in a moving frame of reference, the evolutionary change due to spatial sorting (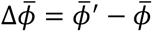; here defined as the mean trait value of offspring in the vacant patches minus the mean trait value of parents in the occupied patches before dispersal) is given by

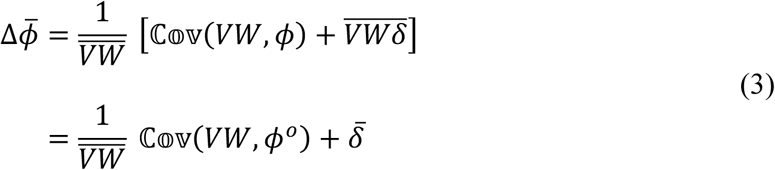

Assuming that the joint distribution of parent and offspring trait values is a multivariate normal distribution, the mean trait value of offspring of the 𝑖th parent can be expressed as

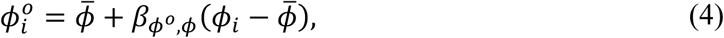

where 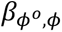 is the slope of the regression between mean offspring 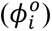 and parents’(𝜙_𝑖_) trait values (Rice 2004). This regression coefficient, 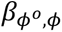, is commonly referred to as heritability (ℎ^2^) of the trait and is equal to the ratio of the additive genetic variance 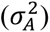 and phenotypic variance 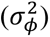 in the parent population (Falconer 1960, Walsh and Lynch 2018). Substituting equation (4) in equation (3) yields an evolutionary change of

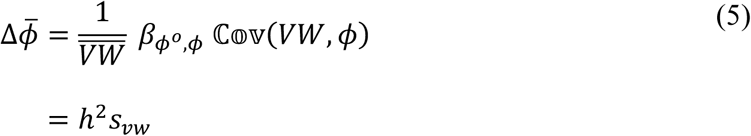

where 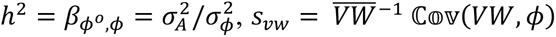, and 𝛿̅ = 0 (See Appendix for details). Alternatively, using the identity for covariance, equation (5) can be re-expressed as

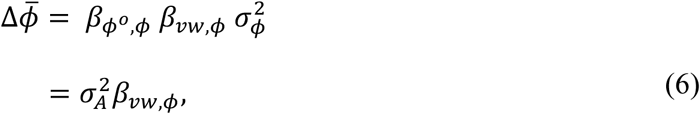

where 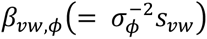 is the slope of the regression line between 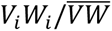 and 𝜙_𝑖_. Intuitively, equation (6) suggests that, in a moving frame of reference, the magnitude of evolutionary change is governed by the heritable variation in the trait and the strength of the statistical association between the number of offspring a parent leaves in the vacant patches and its trait value. This association arises because parents with traits that confer them a dispersal or reproductive advantage are likely to leave more offspring in the vacant patches. This finding highlights the key difference between spatial sorting and natural selection: variation in dispersal propensity can lead to evolution by spatial sorting, even in the absence of differential lifetime reproductive success (Shine et al. 2011; see Figs. S1 and S2 for a more detailed discussion).

Note that the equation of evolution by spatial sorting (Eq. 5) is structurally similar to the Breeder’s equation (Phillips and Perkins 2019). Consequently, we adopt the interpretation and the naming convention of variables in equations (5) and (6) that are analogs to their counterpart variables in the Breeder’s equation. We refer to 𝑠_𝑣w_ as the *sorting differential*, 𝛽_𝑣w,𝜙_ as the *sorting gradient*, 𝑉_𝑖_𝑊_𝑖_—the number of offspring the 𝑖th parent leaves in the vacant patches—as the *sorting fitness* of 𝑖, 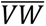 as the *mean population sorting fitness*, and 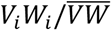 as the *relative sorting fitness* of 𝑖.

More importantly, the similarities between the two models offer insights into what measure to use to quantify the strength of spatial sorting. We propose that the strength of spatial sorting should be measured using the sorting gradient of the standardized phenotypic trait, which we refer to as the *standardized sorting gradient*—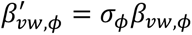. In the case of a single phenotypic trait, 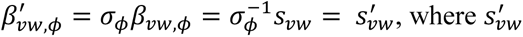 is the *standardized sorting differential*. Using the standardized sorting gradient to quantify the strength of spatial sorting offers several advantages. First, the sorting gradient is closely linked to the equation of evolutionary change by spatial sorting (Eq. 5) and, therefore, has predictive value in the context of quantifying evolutionary changes in phenotypic traits. Second, standardizing phenotypic traits provides a measure of the strength of spatial sorting, which can be compared across biological systems by putting traits on a common scale (Matsumura et al. 2012). Third, due to the structural congruence between the equations of evolution of quantitative traits by natural selection and spatial sorting, standardized sorting gradients can be compared with standardized selection gradients to evaluate the relative magnitude of the two evolutionary forces in generating phenotypic shifts.

## Data Analysis

To demonstrate the practical utility of our model in quantifying the strength of spatial sorting and corresponding evolutionary change, we analyze an exemplar of spatial sorting—survey data on soapberry bugs recolonizing habitats that were extirpated by a hurricane (Comerford et al. 2023).

### Soapberry Bugs

The soapberry bugs’ ancestral range in the USA falls along the southern U.S. border with Mexico and the states abutting the Gulf Coast (Tsai et al. 2013). In humid tropical regions, including the Greater Houston area, soapberry bugs breed year-round. Upon mating, the females lay eggs in 1 cm deep holes beneath their host plants of the soapberry family (Carroll 1988). It takes about 10 days before the larval nymphs emerge from the eggs. Subsequently, the nymphs undergo five molting stages (also referred to as instars) over the next 29 ± 4 days before reaching sexual maturity. These newly molted adult females avoid mating for 6 ± 1 days, after which they lay eggs throughout most of their 60-day lifespan (Carroll 1991) (see Fig. S3).

Female soapberry bugs generally tend to begin histolysis of flight muscles during oogenesis (Carroll et al. 2003), and males show prolonged mate guarding behavior (Carroll 1988). As a result, the potential movement of females between habitats after mating is significantly reduced (Comerford et al. 2023). These insects have binary wing morphologies—long-winged ‘macropterous’ and short-winged ‘brachypterous’ forms (Fig. 1A) (Dingle and Winchell 1997). These binary forms are strongly associated with dispersal abilities of both male and female insects—macropterous insects have fully developed wings that allow them to fly long distances, while flightless brachypterous insects lack flight muscles. Previous studies suggest that this discrete wing polyphenism is determined by juvenile hormones, which, in turn, are regulated by multiple genes with small effects, in addition to resource availability in the environment (Dingle and Winchell 1997, Dingle 2001). Due to life history tradeoffs, wing morphologies also determine variation in fitness of these insects. The high energetic cost of developing and maintaining long wings and flight muscles in macropterous individuals requires longer developmental periods and reduced lifetime fecundity (Zera and Denno 1997). In contrast, brachypterous individuals have higher lifetime fecundity.

**Figure 1:**
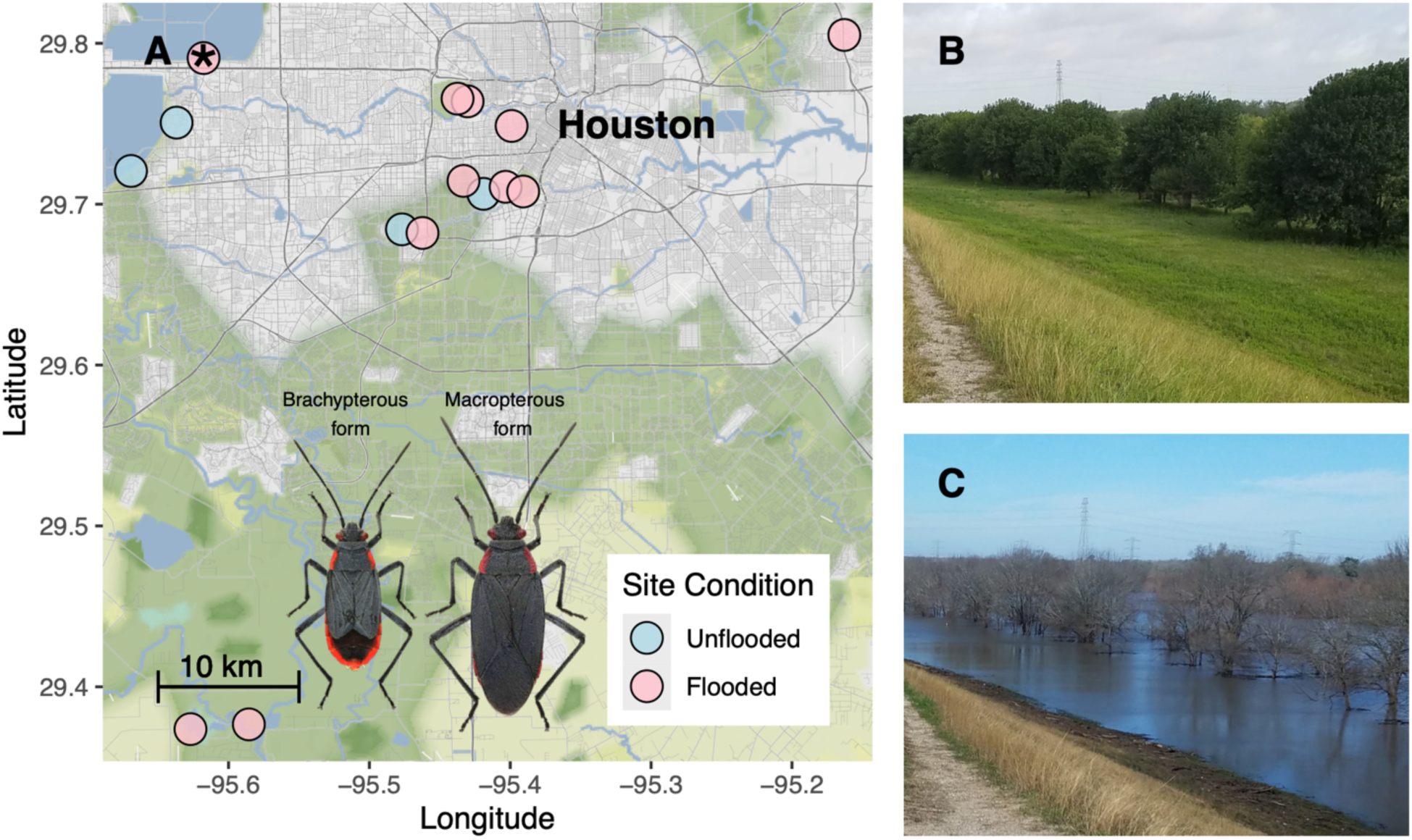
Map of survey sites across the Greater Houston area in Texas where Comerford et al. (2023) sampled soapberry bugs (**A**). Out of the fifteen sampling sites, eleven (pink) experienced local extinction due to prolonged flooding, while four sites (blue) maintained a stable population throughout the survey period. The photographs on the right show a flooded site (marked with an asterisk in **A**) before (**B**) and after (**C**) Hurricane Harvey from the same vantage point. The image of the macropterous and brachypterous forms of the soapberry bug in **A** was taken from Tsai et al. (2013). Photography credit: Mattheau Comerford.

### Hurricane Harvey

Warm sea surface temperatures are perfect incubators for tropical storms in the Gulf of Mexico (Chen et al. 2008, Risser and Wehner 2017). On August 25, 2017, Hurricane Harvey struck southeastern Texas, releasing in excess of 1,500 mm of precipitation in a span of seven days (Trenberth et al. 2018). This one-in-ten-thousand-year event led to intense flooding across the Greater Houston area (Emanuel 2017, Van Oldenborgh et al. 2017). However, not all regions were affected equally. Variation in ground porosity and subsidence led to prolonged flooding in some areas, while others remained unaffected (Miller and Shirzaei 2019). Fortuitously, the hurricane occurred during a multi-year population survey by Comerford et al. (2023), which was initiated a year before the hurricane and continued for an additional three years.

### Monitoring Scheme

Comerford et al. (2023) noted the insect abundances and the morphological form of soapberry bugs every 2 weeks at fifteen sites across the region (Fig. 1A). The survey data revealed two important observations. First, four of the fifteen sites that did not experience prolonged flooding (hereafter called unflooded sites) maintained a stable population throughout the survey (blue points in Fig. 1A). The remaining eleven flooded sites (pink points in Fig. 1A) experienced a local extinction of insects (Fig. 1 B and C). Extinction was confirmed by a gap of 4.5 to 27 months in insect detection at the flooded sites, exceeding the ∼100-day life expectancy of soapberry bugs (from the egg stage to the death of an insect). In addition, inundation experiments by Comerford et al. (2023) showed that after a week of submergence, egg survival drops precipitously and is zero after ten days. This provided further corroborative evidence for local extinction because females lay eggs in the soil beneath the host plants (Carroll 1988), and the topsoil at the flooded sites was inundated for 7 to 90+ days after the hurricane (Fig. 1C). Second, before the hurricane, two thirds of sampled male and female individuals in the unflooded sites were macropterous insects (Fig 2A). In contrast, during the six weeks after the hurricane, starting from the first co-detection of male and female insects, the survey showed that the flooded sites were re-established only by macropterous male and female insects (a total of 88 individuals were sampled). We chose this specific sampling time period because it takes about six weeks for an egg to develop into a penultimate nymph (the last stage before the insect reaches reproductive maturity). During this development period, instars can be visually distinguished from adults, allowing us to avoid individuals that were born in flooded sites after recolonization (see Fig. S3). These survey observations are consistent with the theory of spatial sorting—since macropterous insects are better dispersers than flightless brachypterous insects, insects recolonizing the flooded sites have a higher proportion of macropterous forms relative to the insect population in the unflooded sites before recolonization.

**Figure 2:**
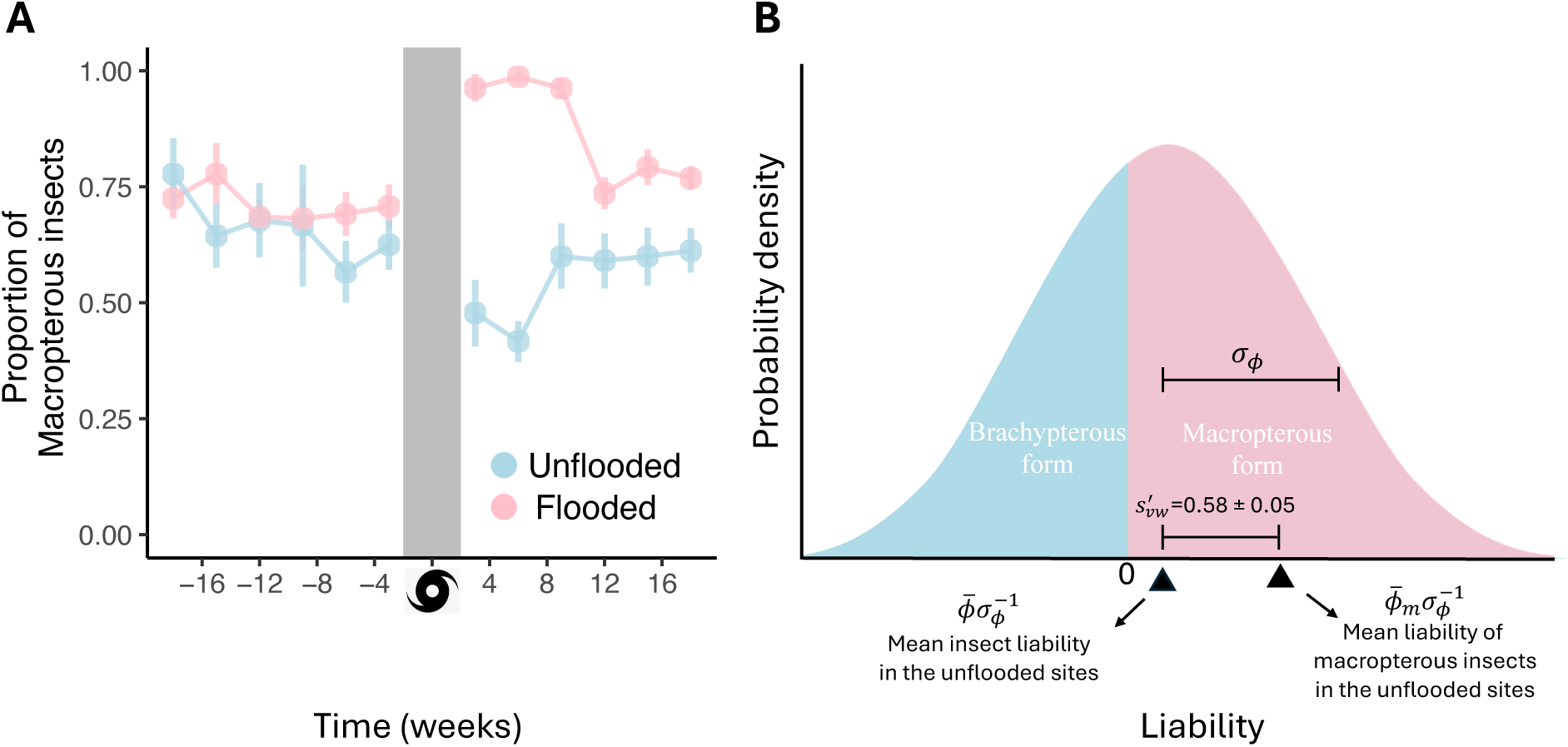
(**A**) The survey data show that, pre-hurricane, both flooded and unflooded sites have a similar proportion (± 1 sd.) of macropterous insects. However, post-hurricane, the initial colonizers in the flooded sites (six weeks after the first detection) had macropterous form, suggesting a non-random colonization event. We also observe a decline in the proportion of macropterous insects in unflooded sites, potentially due to the emigration of macropterous insects. Ten weeks post-hurricane, as the density of insects increases, the proportion of brachypterous insects in the flooded sites approaches the pre-hurricane level, thereby eroding the signature of spatial sorting over time. (**B**) A normal liability distribution of insects in the unflooded sites before the hurricane assumes that insects with liability above zero take macropterous form, such that the area corresponding to the pink region is equal to the proportion of macropterous insects (𝑝). We find that the mean liability of macropterous insects recolonizing the flooded sites (right triangle; mean of the pink region of the liability distribution) is 0.58 ± 0.05 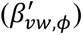 standard deviation higher than that of insects in unflooded sites before the hurricane (left triangle; mean of the full liability distribution). The estimated standardized sorting gradient 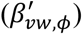 is in the top 10 percentile of standardized selection gradients observed in natural populations (Kingsolver et al. 2001). Genetic crosses show that the liability is a heritable trait (ℎ^2^ = 0.65 ± 0.13). Consequently, spatial sorting resulted in a standardized evolutionary change of 0.37 ± 0.08. The variation in the parameter estimates corresponds to the standard deviation of the posterior distribution. For details on the statistical analysis, refer to the supplementary R codes.

### Strength of Spatial Sorting

To calculate the strength of spatial sorting, we assume the visible morphological form of an insect has a threshold response to an underlying unobserved continuous trait, commonly referred to as liability (Falconer 1960), such that when the liability of an insect is above (below) zero, the individual takes the macropterous (brachypterous) form (Fig. 2B; see Roff (1996) for discussion on thereshold models to describe evolution of disrcrete traits that are regulated by an underlying continious character). Based on this assumption, the proportion of macropterous insects in a population is simply the area under the liability distribution from zero to infinity. The variation in liability is assumed to be determined by environmental and polygenic factors and can be thought of as the concentration of a substance, such as juvenile hormones (Dingle and Winchell 1997, Dingle 2001).

Based on our derivation, the standardized sorting gradient can be expressed as

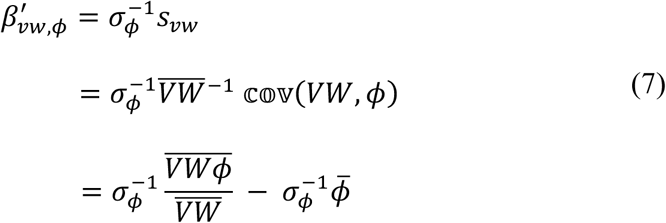

Because the survey data revealed that only macropterous insects participated in the recolonization of flooded sites, we set 𝑉_𝑖_ to zero and a finite value for brachypterous and macropterous insects, respectively. Furthermore, based on the life-history of these insects, we assumed that the fecundity, 𝑊_𝑖_, of the brachypterous insects is greater than that of the macropterous insects. These assumptions allowed us to simplify equation (7) as

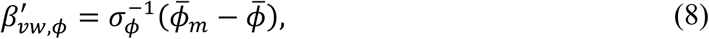

where *φ̄* and *φ̄*_*m*_ are the mean liabilities of all insects and macropterous insects in the unflooded sites before recolonization, respectively (see Appendix for details).

Next, assuming the liability of insects has a normal distribution (Fig. 2B), the mean liability of insects in the unflooded sites (in units of 𝜎_𝜙_) is

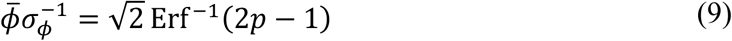

and the mean liability of macropterous insects in the unflooded sites (in units of 𝜎_𝜙_) is

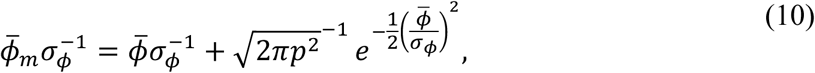

where Erf^-1^ is the inverse error function and 𝑝 is the proportion of macropterous insects in the unflooded sites before recolonization. Intuitively, 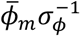 corresponds to the mean of the liability distribution truncated from below at zero (expectation of the pink region of the liability distribution in Fig. 2B). By rearranging the terms in equations (8-10), we can show that the standardized sorting gradient is given by

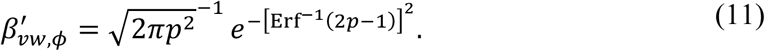

To estimate 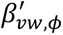, we took six weeks of pre-hurricane data from unflooded sites (Comerford et al. 2023). The survey revealed that 180 of the 280 sampled individuals at the unflooded sites had macropterous forms. Since the insects were sampled randomly, we statistically modeled the proportion of macropterous insects at the unflooded sites (𝑝) as

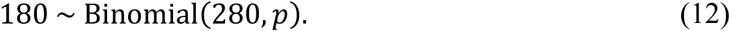

Using the posterior samples of 𝑝 as input in equation (11), we obtained the posterior distribution of 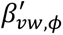 (Carpenter et al. 2017). We found 𝑝 = 0.64 ± 0.03 and 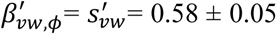. Here, the first and second terms in the estimate of 𝑝 and 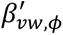 correspond to the mean and standard deviation of the posterior distribution, respectively (see supplementary code for statistical analysis). When we compared the magnitude of the standardized sorting gradient to the magnitude of standardized selection gradients documented in the scientific literature, it ranked in the top ten percentile among ∼1400 previous studies (Kingsolver et al. 2001).

### Heritability and Evolutionary Response to Spatial Sorting

Due to non-destructive sampling, the survey efforts by Comerford et al. (2023) were unable to distinguish macropterous immigrants from macropterous offspring born in the flooded sites. As a result, we could not estimate proportion of macropterous offspring in the flooded sites from survey data, which is required to estimate the realized evolutionary change (see Eq. 2). To address this challenge, we performed multigenerational crosses to estimate heritability (see Supplementary Information for details), which was subsequently used as input in our quantitative genetics model (Eq. 5) to estimate the evolutionary response. This multigenerational breeding experiment was conducted under standardized laboratory conditions, which allowed us to control for plasticity and maternal effects.

We collected 40 juvenile insects from three geographical locations (Austin, Stock Island, and Gulfport), which are separated by more than 350 km. The insects were reared under standardized laboratory conditions and randomly mated to generate 28 males and 28 females of the F1 generation of insects. Upon maturation, the morphological form of F1 insects was noted: we used the numerical value one (zero) to represent macropterous (brachypterous) insects. Subsequently, the insects were randomly paired to generate an F2 generation of insects. Upon eclosion from eggs, 14 F2 nymphs were selected haphazardly from each maternal line. The insects were reared under the same conditions as F1 insects, and their morphological form was noted upon maturation.

To estimate heritability, we fitted a Bayesian animal model (De Villemereuil et al. 2016) using the morphology of F1 and F2 insects as input for parents and offspring, respectively. We found a statistically significant estimate of heritability of liability (ℎ^2^ = 0.65 ± 0.13) (Hadfield 2010), suggesting that wing morphology is highly heritable. Finally, we used the posterior samples of ℎ^2^ and 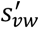 to calculate the posterior distribution of standardized evolutionary change 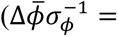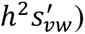. We find that non-random recolonization of the flooded sites would have resulted in a standardized evolutionary change of 0.37 ± 0.08 in insect liability. The large magnitude of standardized evolutionary change suggests that evolution by spatial sorting may exert a strong influence on the spatial population dynamics of a species.

## Discussion

We present a quantitative genetics model of spatial sorting to describe evolution in a patchy landscape (Eq. 5). We find that, in a moving frame of reference, evolution by spatial sorting is driven by heritable variation in a trait that confers differential sorting fitness—the number of offspring a parent leaves in the vacant patches. We show that the evolutionary change can be partitioned into two components, sorting gradient and additive genetic variance, which can be estimated independently using traits of organisms measured during a recolonization event and an animal model with pedigree data as input, respectively. We propose that, analogous to the Breeder’s equation, the sorting gradient of standardized traits (i.e., 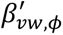) can be used to quantify the strength of spatial sorting. This choice offers two advantages. First, putting traits on a common scale (*i.e.*, in units of standardized deviation) allows evolutionary biologists to compare the strength of spatial sorting across biological systems (Matsumura et al. 2012). Second, because of the structural congruence between the quantitative genetics model of spatial sorting and the Breeder’s equation, biologists can juxtapose standardized sorting and selection gradients to assess the relative magnitude of phenotypic shifts arising from spatial sorting and natural selection.

We applied our quantitative genetics model of spatial sorting to study the evolution of soapberry bugs’ liability because of morphology-dependent recolonization of habitats extirpated by Hurricane Harvey in Greater Houston (Fig. 1; Comerford et al. 2023). We found a statistically significant standardized sorting gradient 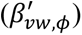 of 0.58 ± 0.05 (Fig. 2), which ranked in the top ten percentile of standardized selection gradients documented in the scientific literature (Kingsolver et al. 2001). Genetic crosses revealed that insect liability is a heritable trait (ℎ^2^ = 0.65 ± 0.13), yielding a standardized evolutionary change in liability 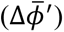 of 0.37 ± 0.08 due to spatial sorting.

The estimate of the standardized sorting gradient suggests that non-random colonization of habitats can produce phenotypic shifts comparable to the most extreme observed phenotypic shifts in natural populations (Kingsolver et al. 2001). And, since many natural populations have abundant heritable variation in dispersal-associated traits (Saastamoinen et al. 2018), spatial sorting may yield evolutionary changes on ecological timescales, which can, in turn, produce novel eco-evolutionary feedback (Miller et al. 2020).

Because of the structural similarities between the Breeder’s equation and the quantitative genetics model presented in the paper, our model can be extended to incorporate additional features, such as genetic correlations between multiple traits (Lande and Arnold 1983) arising from pleiotropy and linkage. These correlations can dramatically change inferences about the direction and the magnitude of spatial sorting obtained from univariate analysis. For example, conducting separate univariate analyses on two strongly correlated traits with the same signed phenotypic shifts will yield the same signed sorting gradients. However, a joint multivariate analysis may reveal that the traits may have opposite-signed sorting gradients, even though the observed phenotypic shifts after spatial sorting have the same sign. This situation can arise when the indirect response to spatial sorting due to genetic correlations on one of the traits is much larger in magnitude than the direct response.

Genetic correlations can also be important when there is a sex-specific response to spatial sorting (Lande 1980, McGlothlin et al. 2019). For example, when mating precedes dispersal, the probability of leaving offspring in the vacant patches depends mainly on the phenotype of females. Consequently, the evolutionary response in males will be dominated by an indirect response to spatial sorting due to the shared genome between the two sexes.

Our theoretical framework also has several practical limitations. The model describes evolution over a single generation in a patchy landscape with local extinction and colonization dynamics. Although spatial sorting has been documented in many patchily distributed populations (Hill et al. 1999, Hanski et al. 2004, Forsman et al. 2011), the model may not be appropriate to describe evolution in spatially structured populations, such as invasion fronts (Phillips et al. 2006) and range expansions (Duckworth and Badyaev 2007), where phenotypic changes due to spatial sorting can compound over multiple generations and density dependent effects may play an important role (Shaw et al. 2023). This is because our model assumes that, during spatial sorting, individuals disperse to empty habitats and ignores feedback between ecological and evolutionary dynamics, which makes the model formulation analytically intuitive and simplifies the estimation of sorting gradients using phenotypic data from patchy populations. However, these assumptions are unlikely to hold at population fronts, complicating theoretical modeling of evolutionary dynamics and quantification of the sorting gradient in these natural settings. Theoretical extension of our model, which integrates population dynamics with a quantitative genetics model of spatial sorting (see Pease et al. 1989, as an example of how to integrate Breeder’s equation with a spatial population dynamics model), may offer new insights into how density-dependent effects shape trait clines.

Nevertheless, in some special cases, the phenotypic analysis presented in the paper can also be applied to structured populations. For example, in early stages of population expansion or when a new satellite colony is formed far from the invasion front, a biogeographer can divide the landscape into a well-mixed core habitat and a periphery region. The core (periphery) region can be treated as occupied (unoccupied) patches in our model. Now, by tracking the movement and phenotypes of individuals from the core (thorough mark and recapture or animal trackers) to the periphery, one can estimate the sorting fitness, which can, in turn, be used to estimate the sorting gradient.

To summarize, our quantitative genetics model and empirical analysis provide quantitative evidence that, like natural selection, spatial sorting can produce phenotypic shifts on ecological timescales. As a result, spatial sorting is likely to play an important role in the Anthropocene in determining the distribution of organisms and spatial genetic variation. This general approach will serve as the foundation for additional work quantifying the magnitude of spatial sorting under different scenarios in an era of global change, where movement is becoming increasingly important.

## Acknowledgment

Nikunj Goel was supported by the Continuing and Stengl-Wyer fellowships by the University of Texas at Austin, Timothy Keitt was supported by the Planet Texas 2050 grant from the University of Texas at Austin. The monitoring survey of soapberry bugs was funded by the National Science Foundation to Scott P. Egan (Award Number: 1802715) and NSF-PRFB (Award Number: 2305864) to Mattheau S. Comerford. We thank Robert Heckman for his help with setting up the animal model. We also thank Dan Bolnick, Mevin Hooten, and Xinyi Yan for providing valuable feedback, and Dr. Scott Carroll for introducing Mattheau S. Comerford and Scott P. Egan to the soapberry bug system.

## Author Contributions

NG conceived and designed the study based on feedback from TEJ, MSC, and THK. MSC led the survey and performed the crosses with feedback from SPE. NG built the model and analyzed the data with feedback from TEJ. NG wrote the paper with feedback from all authors. Authors declare no conflict of interest.

