## Supplementary Information for "Measuring the strength of spatial sorting"

### Calculating $\Delta\bar{\phi}$

In the main text, we derived

$$\Delta\bar{\phi} = \frac{1}{\overline{VW}} \mathbb{Cov}(VW, \phi^o) + \bar{\delta}.$$

To simply the above equation, note that

$$\bar{\delta} = N^{-1} \sum_{i=1}^{N} \delta_i$$

$$= N^{-1} \sum_{i=1}^{N} (\phi_i^o - \phi_i)$$

Substituting equation (4) in the above expression, we get

$$\bar{\delta} = \left(\beta_{\phi^o,\phi} - 1\right) N^{-1} \sum_{i=1}^{N} (\phi_i - \bar{\phi})$$

$$= \left(\beta_{\phi^o,\phi} - 1\right)(\bar{\phi} - \bar{\phi}) = 0.$$

This yields

$$\Delta\bar{\phi} = \frac{1}{\overline{VW}} \mathbb{Cov}(VW, \phi^o)$$

Intuitively, $\bar{\delta} = 0$ implies that, in the absence of differential dispersal and reproduction, the mean trait value does not change (see pages 178 and 179 in Rice 2004). Now, we again substitute equation (4) in the above equation

$$\Delta\bar{\phi} = \frac{1}{\overline{VW}} \mathbb{Cov}\left(VW, \bar{\phi} + \beta_{\phi^o,\phi}(\phi - \bar{\phi})\right)$$

$$= \frac{1}{\overline{VW}} \mathbb{Cov}\left(VW, \beta_{\phi^o,\phi}\phi\right)$$

$$= \frac{1}{\overline{VW}} \beta_{\phi^o,\phi} \mathbb{Cov}(VW, \phi).$$

**Calculating $\overline{VW\phi}/\overline{VW}$**

Let's us assume that there are $N_m$ and $N_b$ macropterous and brachypterous insects in the unflooded patches, respectively, such that $N = N_m + N_b$. Assume the fitness of macropterous and brachypterous insects is $W_m$ and $W_b$, and $V_m$ and $V_b$ are their respective dispersal probabilities. Based on the monitoring data and life-history tradeoffs, we set $V_m \neq 0$, $V_b = 0$, and $W_b > W_m$. We can show

$$\frac{\overline{VW\phi}}{\overline{VW}} = \frac{\sum_{i=1}^{N} V_i W_i \phi_i}{\sum_{i=1}^{N} V_i W_i}$$

$$= \frac{\sum_{i=1}^{Nm} V_m W_m \phi_i^m + \sum_{i=1}^{Nb} V_b W_b \phi_i^b}{\sum_{i=1}^{N_m} V_m W_m + \sum_{i=1}^{N_b} V_b W_b},$$

where $\phi_i^m$ and $\phi_i^b$ is the liability of the $i$th macropterous and brachypterous insect in the unflooded sites, respectively. Since $V_b = 0$, we get a simplified expression

$$\frac{\overline{VW\phi}}{\overline{VW}} = \frac{\sum_{i=1}^{N_m} V_m W_m \phi_i^m}{\sum_{i=1}^{N_m} V_m W_m}$$

$$= \frac{\sum_{i=1}^{N_m} \phi_i^m}{N_m} = \bar{\phi}_m,$$

where $\bar{\phi}_m$ is the mean liability of macropterous insects in the unflooded patches before the hurricane. The above result implicitly assumes that among macropterous insects, the liability of insects does not covary with the sorting fitness, i.e., $\mathbb{Cov}(V_m W_m, \phi_m) = 0$.

**Genetic crosses**

To estimate the heritability of insects' liability, we collect 40 wildtype juvenile soapberry bugs from three geographically separated populations located in the southern United States: Stock Island (24°34'27.1"N 81°44'54.8" W), Gulfport (27°44'56.8"N 82°42'22.7" W) and Austin (30° 12' 59.76"N 97° 39' 2.51" W). Once captured, the juvenile bugs were transported back to the laboratory and housed by deme in three 34.6 x 21 x 12.1cm plastic cages, where they were mass-reared to generate an additional generation (F1) of insects to minimize maternal effects. To control for plasticity, all insects were reared in an environmental chamber held at 29˚ C with 47% RH under a 14L:10D photoperiod to best match the average ambient conditions of all three collection locations. Each rearing container was lined with 6mm of sterile sand to provide females with adequate oviposition substrate. Likewise, a 15 x 15cm sheet of plastic chicken wire mesh was placed in each container as an additional climbing surface. Insects in all three demes were fed *ad libitum* on the seeds of their ancestral host plant balloon vine (*Cardiospermum halicacabum*) purchased from Outsidepride Seed Source LLC (Salem, OR) and provided water from cotton stopped 1.5mm vials.

We randomly selected unmated 28 F1 males and 28 F1 females from within and between populations. We placed them as mate pairs in separate 473ml plastic deli cups, which were lined with sand at the bottom and a mesh cover at the top for ventilation. One week after the mating occurred, F1 insects were sacrificed, pinned, and their morphological form (macropterous vs brachypterous) was noted. Upon eclosion from eggs, 14 F2 neonatal nymphs were selected haphazardly from each maternal line, and the remaining nymphs were sacrificed and discarded. Upon reaching adulthood, F2 individuals were sacrificed, pinned, and their morphological form was noted.

The information gathered from the experiment was collated into two parts. The first part was the pedigree relationships describing the genetic relationship between F1 and F2 insects. The second part was the morphological form of F1 and F2 insects. We represented 'macropterous' insects as one and 'brachypterous' insects as zero. Finally, using MCMCglmm (De Villemereuil et al. 2016), with pedigree and morphology data as inputs, we fit a threshold model (with probit link function) to obtained the posterior distribution of heritability of liability ($h^2 = 0.65 \pm 0.13$).

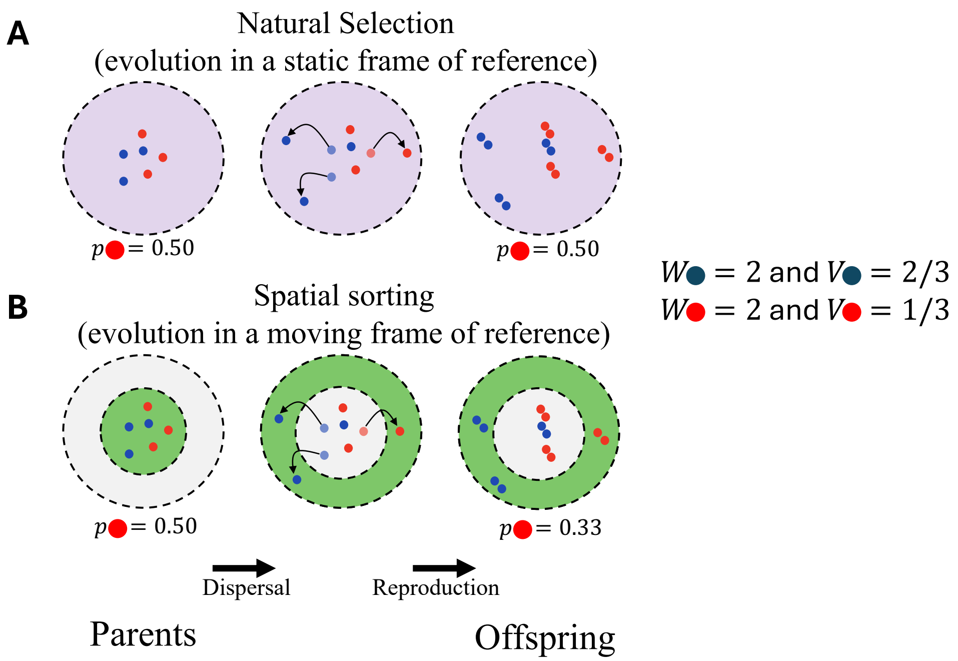


**Figure S1:** Quantifying evolutionary change in **(A)** a static frame of reference (natural selection) and **(B)** a moving frame of reference (spatial sorting) when the red and blue individuals have the same lifetime reproductive success but differ in their dispersal ability. In both **(A)** and **(B)**, parents (left column) follow the same life cycle: disperse, produce offspring (right column), and then die. **(A)** As expected, when evolution is measured in a static frame of reference (frequency of red offspring in the purple region minus the frequency of red parents in the purple region), we do not see a change in the frequency of blue and red types due to a lack of differences in fitness. **(B)** However, when evolution is measured in a moving frame of reference (frequency of red offspring in the green region minus the frequency of red parents in the green region), we see an increase in the frequency of blue individuals at the vanguard. This simple illustration shows that, in a moving frame of reference, variation in dispersal capacity can lead to evolution by spatial sorting, even in the absence of differential lifetime reproductive success.

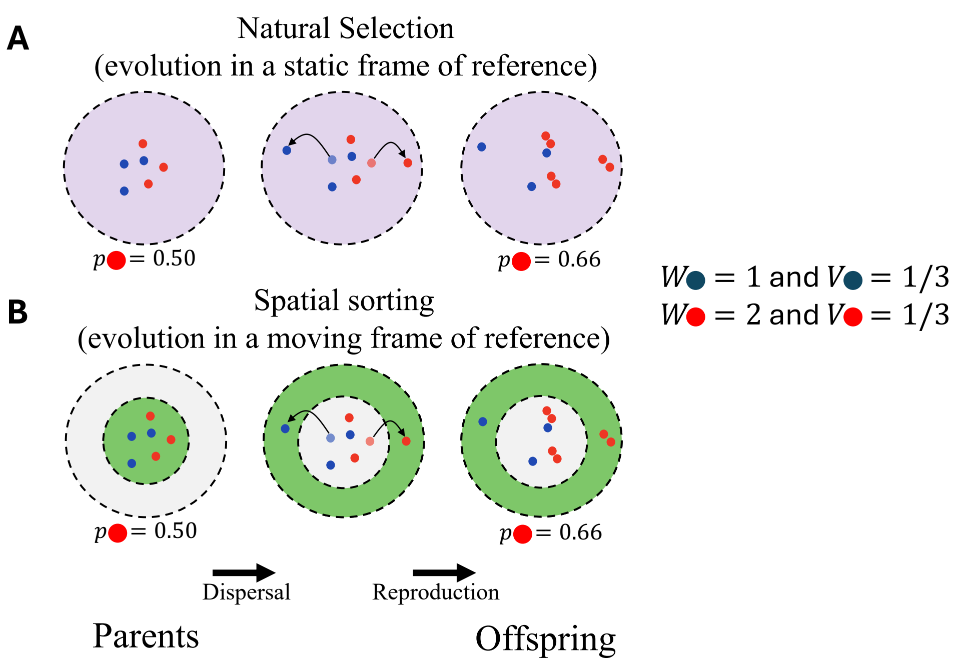


**Figure S2:** Quantifying evolutionary change in **(A)** a static frame of reference (natural selection) and **(B)** a moving frame of reference (spatial sorting) when the red and blue individuals have the same dispersal capacity but differ in their lifetime reproductive success. In both **(A)** and **(B)**, parents (left column) follow the same life cycle: disperse, produce offspring (right column), and then die. **(A)** As expected, when evolution is measured in a static frame of reference (frequency of red offspring in the purple region minus the frequency of red parents in the purple region), we see an increase in the frequency of red individuals due to differences in fitness of red and blue individuals. **(B)** When evolution is measured in a moving frame of reference (frequency of red offspring in the green region minus the frequency of red parents in the green region), we see an increase in the frequency of red individuals at the vanguard. This illustration shows that, although not necessary, differential lifetime reproductive success, too, can lead to evolution by spatial sorting. Thus, both differential dispersal capacity and differential lifetime reproductive success (i.e., differential sorting fitness) can lead to evolution by spatial sorting.

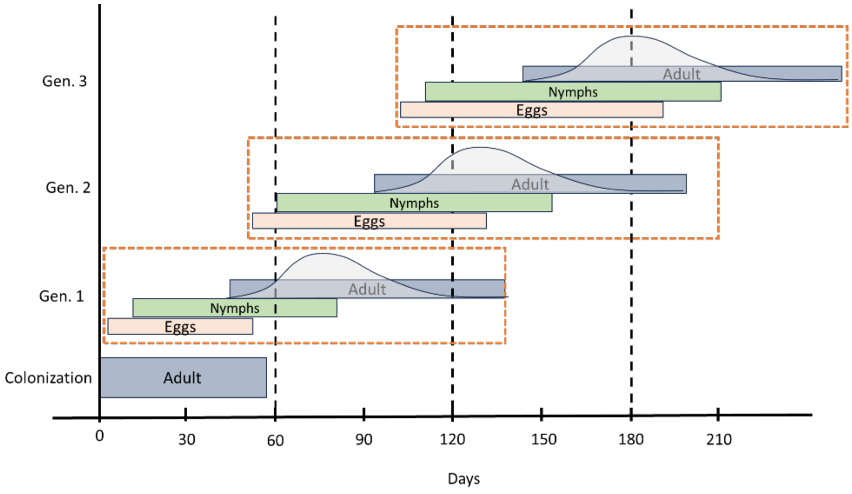


**Figure S3:** The plot shows the development stages of insects arriving at the flooded sites. After females arrive at the flooded sites, they mate with males, upon which female flight muscles begin to histolyze (Carroll 1988). Females lay eggs throughout their adult life span. After 10 days, larval nymphs emerge from the eggs (Carroll 1991). Subsequently, they undergo four molting stages over 29 ± 4 days before reaching reproductive maturity (fifth instar) (Tsai et al. 2013). The first four instar stages can be visually distinguished from the fifth instar. Based on the life-history cycle of the soapberry bugs, we assume that the fifth instar insects observed at the flooded site until six weeks from the first co-detection male and female insects were not born in the flooded sites.
